# Delayed maturation of the structural brain connectome in neonates with congenital heart disease

**DOI:** 10.1101/2020.09.21.306084

**Authors:** Maria Feldmann, Ting Guo, Steven P. Miller, Walter Knirsch, Raimund Kottke, Cornelia Hagmann, Beatrice Latal, Andras Jakab

**Affiliations:** Child Development Center, University Children‘s Hospital Zurich, Zurich, Switzerland; Children’s Research Center, University Children’s Hospital Zurich, Zurich, Switzerland; Neurosciences and Mental Health, The Hospital for Sick Children Research Institute, Toronto, Canada; Department of Paediatrics, The Hospital for Sick Children and the University of Toronto, Toronto, Canada; Division of Pediatric Cardiology, Pediatric Heart Center, University Children’s Hospital Zurich, Zurich, Switzerland; Department of Diagnostic Imaging, University Children’s Hospital Zurich, Zurich, Switzerland; Department of Neonatology and Pediatric Intensive Care, University Children’s Hospital Zurich, Switzerland; Centre for MR Research, University Children’s Hospital Zurich, Zurich, Switzerland

**Author notes:** Corresponding author: Maria Feldmann.

## Abstract

There is emerging evidence for delayed brain development in neonates with congenital heart disease (CHD). We hypothesize that the perioperative development of the structural brain connectome is a proxy to such delays. Therefore, we set out to quantify the alterations and longitudinal pre- to postoperative changes in the connectome in CHD neonates and assess risk factors for disturbed perioperative network development relative to healthy term newborns. In this prospective cohort study, 114 term neonates with CHD underwent cardiac surgery at the University Children’s Hospital Zurich. Forty-six healthy term newborns were included as controls. Pre- and postoperative structural connectomes were derived from mean fractional anisotropy values of fibre pathways traced using diffusion tractography. Graph theory parameters calculated across a range of proportional cost thresholds were compared between groups by multi-threshold permutation correction adjusting for confounders. Network based statistic was calculated for edgewise network comparison. White matter injury (WMI) volume was quantified on 3D T1-weighted images. Random coefficient mixed models with interaction terms of (i) CHD subtype and (ii) WMI volume with postmenstrual age at MRI respectively were built to assess modifying effects on network development. Pre- and postoperatively, at the global level, efficiency, indicative of network integration, was higher in controls compared to CHD neonates. In contrast, local efficiency and transitivity, indicative of network segregation, were higher in CHD neonates compared to controls (all p<0.025 for one-sided t-tests). Preoperatively these group differences were also found across multiple widespread nodes (all p<0.025, accounting for multiple comparison), whereas postoperatively nodal differences were not evident. At the edge-level, the majority of weaker connections in CHD neonates compared to controls involved interhemispheric connections (66.7% preoperatively; 54.5% postoperatively). A trend showing a more rapid pre- to postoperative decrease in local efficiency was found in class I CHD neonates compared to controls. In CHD neonates, larger WMI volume was associated with lower strength (p=0.0026) and global efficiency (p=0.0097). The maturation of the structural connectome is delayed in neonates with CHD, with a pattern of lower structural integration and higher segregation compared to healthy controls. Trend-level evidence indicated that normalized postoperative cardiac physiology in class I CHD neonates might improve structural network topology. In contrast, the degree of WMI burden negatively impacts network strength and integration. Further research is needed to elucidate how aberrant structural network development in CHD represents neural correlates of later neurodevelopmental impairments.

## Background

Congenital heart disease (CHD) occurs at a prevalence of about 8.2 per 1000 live births in Europe and accounts for one-third of all congenital anomalies (van der Linde *et al*. 2011). More than one-third of all CHD infants require surgical correction or catheter interventions during early childhood, and are considered severe cases of CHD (Ferry 1987). CHD survivors are at an increased risk for a spectrum of neurodevelopmental sequelae and cognitive dysfunction across the lifespan (Karsdorp *et al*. 2007, Huisenga *et al*. 2020).

Numerous studies have described aberrant brain development in CHD neonates, that might underlie the risk for adverse neurodevelopment (Claessens *et al*. 2017, Peyvandi *et al*. 2019). With an onset in the foetal period (Limperopoulos *et al*. 2010), delayed brain development in CHD neonates has been found on the macrostructural, microstructural and metabolic scale (Miller *et al*. 2007, Licht *et al*. 2009, Ortinau *et al*. 2012, von Rhein *et al*. 2015, Claessens *et al*. 2016), with the most severe CHD subtypes being most affected (Limperopoulos *et al*. 2010, Peyvandi *et al*. 2018). Furthermore, it has been suggested that brain dysmaturation is the substrate for white matter injury (WMI), which is the most frequently found brain lesion in CHD neonates (Partridge *et al*. 2006, Dimitropoulos *et al*. 2013, Mulkey *et al*. 2014, Guo *et al*. 2019).

Although a link of aberrant brain development and injury with neuro-developmental outcome has been demonstrated (Andropoulos *et al*. 2014, Claessens *et al*. 2018, Meuwly *et al*. 2019), the complex phenotype of later neurodevelopmental impairments in CHD children remains insufficiently explained. Thus, more advanced neuroimaging analyses considering the complex network of the brain beyond specific structure and function mapping are needed to better capture adverse brain development in CHD neonates (Peyvandi *et al*. 2016).

Structural connectomics is a rapidly emerging tool, that is promising in studying the emergent properties of the global brain network organization (Rubinov *et al*. 2010) and revealing systems-level effects of disease in the developing neonatal brain (Tymofiyeva *et al*. 2013, Song *et al*. 2017, Keunen *et al*. 2018, Jakab 2019). Driven by neurobiological processes during foetal and neonatal development the whole brain network transitions from a highly segregated to a more integrated organisational architecture (Cao *et al*. 2017), allowing for evolving parallel and efficient information processing across remote brain regions (Rubinov *et al*. 2010). As previously shown, the connectome is subject to developmental disturbances in at risk populations, such as infants with neonatal ecephalopathy or after preterm birth (Ziv *et al*. 2013, Pandit *et al*. 2014, Batalle *et al*. 2017). As such, structural connectomics may detect aberrant brain development that appears normal on routine diagnostic imaging.

Recent studies demonstrated aberrations in the architecture of the neonatal whole brain connectome in CHD, indicating possible vulnerability of brain network organization towards developmental disturbances in this population (De Asis-Cruz *et al*. 2018, Schmithorst *et al*. 2018). However, to date the characteristics of the neonatal structural connectome and the development across the perioperative period in CHD neonates in comparison to healthy controls are not well studied. Furthermore, while there is evidence that WMI can disrupt the structural and functional connectome in preterm born infants (Ceschin *et al*. 2015, Cai *et al*. 2017), how the specific emergent properties of the whole-brain network are affected by WMI in neonates with CHD remains poorly understood.

Therefore, the primary aim of this study is to characterise the neonatal structural brain connectome in pre- and postoperative neonates with CHD in comparison to healthy controls. Second, we aim to assess risk factors for adverse network development, such as CHD severity sub-type and WMI. We hypothesize (i) that structural whole brain connectomics will reveal systems-level aberrant brain development beyond what is evident on routine diagnostic imaging; (ii) that there is an effect of CHD subgroups on pre- to postoperative network development and (iii) that WMI is a risk factor for the perturbations of the neonatal connectome.

## Methods

### Study population

In this prospective cohort study neonates with CHD who required neonatal corrective or palliative cardiac surgery during the first weeks of life at the University Children’s Hospital Zurich between December 2009 and March 2019 were eligible. Neonatal corrective or palliative cardiac surgery was considered irrespective of the use of cardiopulmonary bypass surgery. Furthermore, cases with single ventricle physiology who underwent a hybrid approach instead of Norwood I procedure were also included. Exclusion criteria were a suspected or confirmed genetic syndrome or a gestational age below 36 weeks. A comparison group of healthy term born neonates was recruited between January 2011 and April 2019 from the well-baby nursery at the University Hospital Zurich. Inclusion criteria for healthy controls were birth above 36 weeks of gestation and unremarkable postnatal adaptation. None of the healthy controls were admitted to the neonatal ward.

Neonatal, perioperative, surgical and demographic characteristics were prospectively extracted from patient charts. CHD diagnoses were classified into anatomical subclasses according to Clancy et al.: class I, biventricular CHD without aortic arch obstruction; class II, biventricular CHD with aortic arch obstruction; class III, univentricular CHD without aortic arch obstruction; class IV, univentricular CHD with aortic arch obstruction (Clancy *et al*. 2000). Parental written informed consent was obtained, and the study was approved by the ethical committee of the Kanton Zurich, Switzerland (KEK StV-23/619/04). The study was carried out in accordance with the principles enunciated in the Declaration of Helsinki and the guidelines of Good Clinical Practice. This is a convenience sample including all subjects with available neonatal diffusion tensor imaging data acquired during the study period (for more information see flow chart in **Supplementary Fig. 1**). Neuroimaging and neurodevelopmental outcomes in subsamples of this cohort were published previously (Bertholdt *et al*. 2014, Claessens *et al*. 2016, Jakab *et al*. 2019, Meuwly *et al*. 2019).

### Brain MRI

Both CHD neonates and healthy controls underwent brain MRI on a 3.0 T clinical MRI scanner using an 8-channel head coil (GE Signa MR750). CHD neonates were scanned pre- and postoperatively, with serial MRI at both time points if possible. Healthy controls received one postnatal MRI. MRI was acquired during natural sleep with a feed and wrap technique; hearing protection was provided. The scanning protocol included a 3D T1-weighted fast spoiled gradient echo sequence (TR (repetition time)/TE (echo time) 11/5 ms (rounded value, actual value used according to the “minimum setting”), inversion time = 450 ms, flip angle = 12, 1 mm^3^ isotropic). Diffusion tensor imaging (DTI) was acquired using a pulsed gradient spin-echo echo-planar imaging sequence (TR/TE 3950/90.5 ms, field of view =18 cm, matrix = 128×128, slice thickness = 3 mm) with 35 diffusion encoding gradient directions at a b-value of 700 s/mm^2^ and four b = 0 images. During the study period, an upgrade was performed from the HD.xt to the MR750 platform. This was accounted for by including MRI cohort as covariate in all multivariate regression models.

### WMI volume quantification

WMI identified as areas of T1 hyperintensity were manually delineated on 3D T1-weighted images using Display software (http://www.bic.mni.mcgill.ca/software/Display/Display.html) as previously described in detail (Guo *et al*. 2019). Cumulative WMI volume in mm^3^ was defined as lesion volume on either pre- or postoperative MRI or the larger lesion volume in cases where both MRI were affected.

### Diffusion tensor image processing

A custom script was used to process the DTI data by wrapping previously published algorithms as described below (script available upon request). DTI data were visually controlled for artifacts. If repeated sequences were not possible for images with low quality, the data were discarded and excluded (Supplementary Figure 1). Brain masks were created using the BET command in FSL and then revised with morphological operations, as well as manual corrections.

A separate step using topup to correct susceptibility induced distortions was skipped in our study as all B0 images and diffusion-weighted frames were acquired using identical antero-posterior phase encoding directions. Spurious image distortions that originated from a combination of eddy currents and real head movement were corrected with the CUDA 8.0 implementation of the eddy command in the FSL software library (version 6.0) with the following parameters: 4 standard deviations as criterion for classifying a slice as outlier (ol_nstd=4), 10 slice to volume iterations (s2v_niter=10), 6 smoothing iterations, during which a filter width of 10, 6, 4, 2, 0, 0 mm for each subsequent smoothing iterations was used. Next, FA and mean diffusivity (MD) maps were calculated by running the dtifit command in FSL on the motion corrected DTI dataset; tensors were fitted with a weighted least squared algorithm. DTI data were further processed by the Bayesian multi-fiber estimation method BedpostX (CUDA 8.0 implementation for FSL 6.0), estimating up to 3 fiber populations per voxel (Hernandez et al. 2013).

### Anatomical parcellation of regions of interest

Anatomical regions of interest (ROI) were parcellated in the anatomical space of the UNC infant brain atlas for neonates (Shi *et al*. 2011), in which a labelling system corresponding to the Automated Anatomical Labeling (AAL) (Tzourio-Mazoyer *et al*. 2002) was used. In the study, we used 90 cortical and subcortical areas. Next, a custom FA template was created in the UNC space based on the FA images of the control subjects to transfer the UNC-AAL ROIs to the DTI space of each subject. The template generation and non-linear co-registration process has been described in detail previously (Jakab *et al*. 2019).

### Structural connectivity and network construction

Seeds for probabilistic diffusion tractography were defined in native DTI space by keeping the voxels with higher than 50^th^ percentile FA value, determined using whole-brain histogram. Whole-brain (i.e. whole white-matter) tractography was carried out using a probabilistic Runge-Kutta 4^th^ order tractography method implemented in Camino (Parker *et al*. 2003), which used the dyadic vector output of the BedpostX algorithm of FSL. Per voxel, 100 probabilistic samples were tracked, probabilistic nearest-neighbour interpolation was used. A termination criterion of FA <0.1 was used for stopping tractography. Tracts shorter than 15 mm were ignored to reduce the effects of spurious connections between neighbouring cortical ROIs. Undirected structural connectivity networks were generated, where each edge represented the number of probabilistic tracts passing through the white matter and connecting any two ROIs (*conmat* command in Camino). Weighted, undirected connectivity networks were formed. The connectivity or edge weight was defined as the mean fractional anisotropy (FA_mean_) value of all tractography streamlines connecting the regions at the two end-points, as these networks have been shown to be most sensitive to microstructural dysmaturation of the white matter in neonates with CHD (Schmithorst *et al*. 2018). Nodes corresponded to the AAL ROIs in subject space.

### Graph theoretical analysis

Network generation and analysis was performed in R Version 3.6.0 (R Core Team 2019) with the packages igraph Version 1.2.4.2 (Csardi *et al*. 2006) and brainGraph Version 2.7.3 (Watson 2019). FA_mean_ networks were thresholded by means of an iterative proportional cost thresholding procedure to filter spurious connections and eliminate the bias of cost for the comparison of network topology (van den Heuvel *et al*. 2017). To ensure that thresholding resulted in the same absolute number of connections across the majority of subjects, we determined the 5^th^ centile of the distribution of network densities. This T_95_ served as upper level of the proportional thresholding procedure, thus ensuring that 95% of the subjects had the same amount of connections at each threshold. Thereby the bias of cost was reduced, while allowing for the inclusion of very sparse networks. Networks were consecutively thresholded over the cost range from 1% to T_95_. Graph theoretical parameters were calculated for all networks at each threshold.

### Graph theory parameters

To quantify the topological organization of brain networks, and specifically depict the main characteristics of network reorganisation during rapid neonatal brain development, parameters of structural integration and segregation were measured. Calculations were performed as implemented in the igraph and brainGraph package in R and as listed in Rubinov and Sporns (Rubinov et al. 2010). As parameter of structural integration network efficiency was calculated. As measure of structural segregation local efficiency and transitivity were assessed. Furthermore, network strength was quantified. A detailed description of the calculated graph parameters can be found in the **Supplementary Methods**.

### Multi-threshold permutation correction

Multi-threshold permutation correction (MTPC) is a comprehensive and sensitive approach for the statistical comparison of network topologies. It takes the trajectory of graph metrics over the whole network cost threshold range into account and thus overcomes the bias of arbitrary threshold selection for network comparison (Drakesmith *et al*. 2015). Pairwise comparison of global and nodal graph parameters between pre- and postoperative CHD neonates and controls was performed by MTPC as implemented in the brainGraph package. The alpha level for statistically significant group differences was set to 0.025 as two one-sided hypothesis tests were performed to determine the direction of group differences. A more detailed description of the MTPC procedure can be found in the **Supplementary Methods**. To control the inflation of the false discovery rate arising from comparing nodal graph theoretical parameters across 90 ROIs, P-values were adjusted with the Benjamini Hochberg procedure (Benjamini *et al*. 1995).

### Network based statistic

Network based statistic (NBS) was performed to compare the strength of each network connection, i.e. at the edge-level, between CHD neonates and controls. For NBS, networks were thresholded at a single cost threshold at T_95_. In brief, in NBS a GLM was specified at each individual network connection and all edges with test statistics above a pre-defined threshold were identified (here P = 0.0005 for two one-sided hypothesis tests to determine the direction of potential group differences). Among those suprathreshold connections any connected components were identified. By means of permutation testing (here 5000 permutations) the P-value controlled for the family wise error rate was determined for each connected component based on its size (Zalesky *et al*. 2010). We used the implementation of NBS in the brainGraph package in R.

### Statistical analysis

Statistical analyses were performed using R Version 3.6.0 (R Core Team 2019). Descriptive statistics for continuous variables were reported as mean and standard deviation or median and interquartile range as appropriate for sample distribution. Categorical variables were reported as proportions.

All statistical models specified within MTPC or NBS analyses included sex, postmenstrual age (PMA) at MRI and scanning cohort as nuisance variables to correct for confounding. The brain networks and nodal MTPC differences were visualized with BrainNet Viewer for Matlab R2014 (Xia *et al*. 2013).

Random coefficient linear mixed effect models from the *nlme* package were built to assess longitudinal growth trajectories of network parameters among CHD subclasses and the effects of WMI on network development. Therefore, a single network threshold at the T_95_ maximum cost threshold was chosen. To account for the longitudinal structure of the data and allow for individual trajectories (Schielzeth *et al*. 2009) the subject identifier and time at scan were included as random effects. All models included sex and scanning cohort as fixed effects. PMA at scan was mean centred and scaled to have a mean of zero and SD of 1. Two main mixed model analyses were conducted. First, to assess network parameter development among CHD sub-classes and controls an interaction term between subclass and PMA at scan was specified. The control group served as the reference group. To test for further interactions between cardiac subclasses the slopes of the subtypes were compared with each other using the *glht* command from the *multcomp* package in R. Second, to assess the effect of WMI on network development an interaction term between cumulative WMI volume and PMA was specified. Non-significant interaction terms were split into non-interacting fixed effects in the final models. As each mixed model was calculated for four global graph theoretical parameters, the family wise error rate was controlled with the Bonferroni correction. The Bonferroni adjusted P-values are given as P_adj_.

To assess the robustness of global group differences between CHD neonates and healthy controls, we excluded all CHD neonates with WMI or arterial ischemic stroke in a post-hoc sensitivity analysis and repeated the MTPC comparison of pre- and postoperative global graph theory parameters.

### Data availability statement

The de-identified data that support the findings of this study will be made available upon reasonable request from the corresponding author.

## Results

### Study population

122 CHD neonates were included in the study. Among those, 114 CHD neonates had DTI sequences with sufficient quality at least at one time point, which allowed for the reconstruction of the structural connectome and the inclusion in further analyses. For further details on subjects excluded for clinical reasons or poor imaging quality see flow chart in **Supplementary Fig. 1**. Ultimately, 79 preoperative and 91 postoperative connectomes (56 cases with both pre- and postoperative connectomes) were included in this analysis. Of 48 included healthy controls, structural connectomes could be generated in 46 (excluded for motion n=2). Cardiac diagnoses were classified as class I in 66/114 (57.9%), class II in 24/114 (21.1%), class III in 7/114 (6.14%) and class IV in 17/114 (14.9%). Demographic and clinical characteristics of the study cohort can be found in **Table 1**.

**Table 1.** Demographic and clinical characteristics of study population. CHD, congenital heart disease; PMA, postmenstrual age; CPB, cardiopulmonary by-pass.

|  | CHD | Controls | P |
| --- | --- | --- | --- |
| n | 114 | 46 |  |
| Gestational age, weeks, mean (SD) | 39.4 (1.3) | 39.5 (1.3) | 0.75 |
| Birth weight, g, median [IQR] | 3300.0 [3000.0, 3670.0] | 3340.0 [3050.0, 3650.0] | 0.67 |
| Head circumference, cm, median [IQR] | 34.5 [34.0, 35.2] | 35.0 [34.0, 36.0] | 0.057 |
| Sex, n (%) | 83 (72.8) | 21 (45.7) | 0.002 |
| Apgar score at 5 min, median [IQR] | 9.0 [8.0, 9.0] | 9.0 [9.0, 9.0] | 0.22 |
| Mechanical ventilation, days, median [IQR] | 3.0 [2.0, 4.0] | NA |  |
| Intensive care unit stay, days, median [IQR] | 6.0 [4.0, 8.0] | NA |  |
| Univentricular CHD, n (%) | 24 (21.1) | NA |  |
| Cyanotic CHD, n (%) | 14 (12.3) | NA |  |
| CPB surgery, n (%) | 91 (79.8) | NA |  |
| Age at surgery, days, median [IQR] | 10.0 [8.0, 13.8] | NA |  |
| Age at MRI, days, median [IQR] |  | 21.0 [16.0, 27.2] |  |
| preoperatively | 7.0 [6.0, 10.0] |  |  |
| postoperatively | 26.0 [21.0, 34.8] |  |  |
| Postmenstrual age at MRI, weeks, median [IQR] |  | 42.3 [41.2, 43.9] |  |
| preoperatively | 40.3 [39.4, 41.6] |  |  |
| postoperatively | 43.4 [42.0, 44.4] |  |  |
| Days pre- to postoperative MRI, median [IQR] | 19 [14.0, 24.0] | NA |  |
| Days surgery/intervention to MRI, median [IQR] | 14 [10.75, 19.25] | NA |  |

### Brain injury and white matter injury burden

Arterial ischemic stroke was observed in 4/79 (5.1%) CHD neonates preoperatively and 4/91 (4.4%) postoperatively. In subjects with serial pre- and postoperative MRI none of the strokes was new on postoperative MRI. All were focal cortical strokes except one case with involvement of the main branch of the middle cerebral artery. WMI was detected in 13/79 (16.5%) CHD neonates on preoperative MRI. Median [IQR] preoperative WMI volume was 57.72 mm^3^ [34.41, 518.59] (missing quantification n=1 due to missing 3D T1). On postoperative MRI WMI was evident in 12/91 (13.2%) CHD neonates with a median volume of 26.30 mm^3^ [16.41, 48.79]. In 1/7 (14.3%) neonates with postoperative WMI and MRI at both scanning time points, WMI was new.

### Less integrated and more segregated structural brain networks in CHD neonates compared to controls

The mean network density (cost) of the non-thresholded connectivity matrices was 0.29 ±0.06 (range 0.13 - 0.41) and the 5th centile of the cost distribution was 0.19. Thus 0.19 was defined as T_95_ and network analyses were performed across the range from 1 to 19% of network cost.

#### Global network topology

Preoperatively, the MTPC based comparison of global network topology between CHD neonates and controls revealed significantly higher global efficiency in controls compared to CHD neonates. The opposite was found for local efficiency and transitivity which were higher in CHD neonates compared to controls. Postoperatively, the same differences were found: Global efficiency was higher, whereas local efficiency and transitivity were lower in controls compared to postoperative CHD neonates (**Table 2; Fig. 1**).

**Table 2.** MTPC results of global network comparison between pre- and postoperative CHD neonates and controls. Significant global level differences between pre- and postoperative CHD neonates and healthy controls as revealed by MTPC. As two one-sided tests were performed to determine the direction of the effects, results are grouped by the contrast “Controls > CHD” or “CHD > Controls” and are significant at the alpha level of 0.025. Amtpc and Acrit denote the results of the overall MTPC comparison across the whole range of thresholds. *Threshold indicates the cost threshold at which the strongest *ß coefficient was observed. Statistical parameters are given for that threshold.

| Parameter | Contrast | *Threshold | *$\beta$ | *SE | *95% CI | *P | Amtpc | Acrit |
| --- | --- | --- | --- | --- | --- | --- | --- | --- |
| **Preoperatively** |  |  |  |  |  |  |  |  |
| Global efficiency | Controls > CHD | 0.14 | 0.0099 | 0.003 | [0.0032; 0.017] | < 0.001 | 0.26 | 0.16 |
| Local efficiency | CHD > Controls | 0.19 | 0.26 | 0.077 | [0.084; 0.44] | < 0.001 | 0.176 | 0.175 |
| Transitivity | CHD > Controls | 0.15 | 0.044 | 0.0095 | [0.022; 0.065] | < 0.001 | 0.42 | 0.18 |
| **Postoperatively** |  |  |  |  |  |  |  |  |
| Global efficiency | Control > CHD | 0.19 | 0.0087 | 0.0027 | [0.0025; 0.015] | < 0.001 | 0.25 | 0.14 |
| Local efficiency | CHD > Control | 0.19 | 0.19 | 0.065 | [0.044; 0.34] | 0.0019 | 0.44 | 0.19 |
| Transitivity | CHD > Control | 0.16 | 0.034 | 0.01 | [0.011; 0.057] | < 0.001 | 0.23 | 0.17 |

**Fig. 1.**
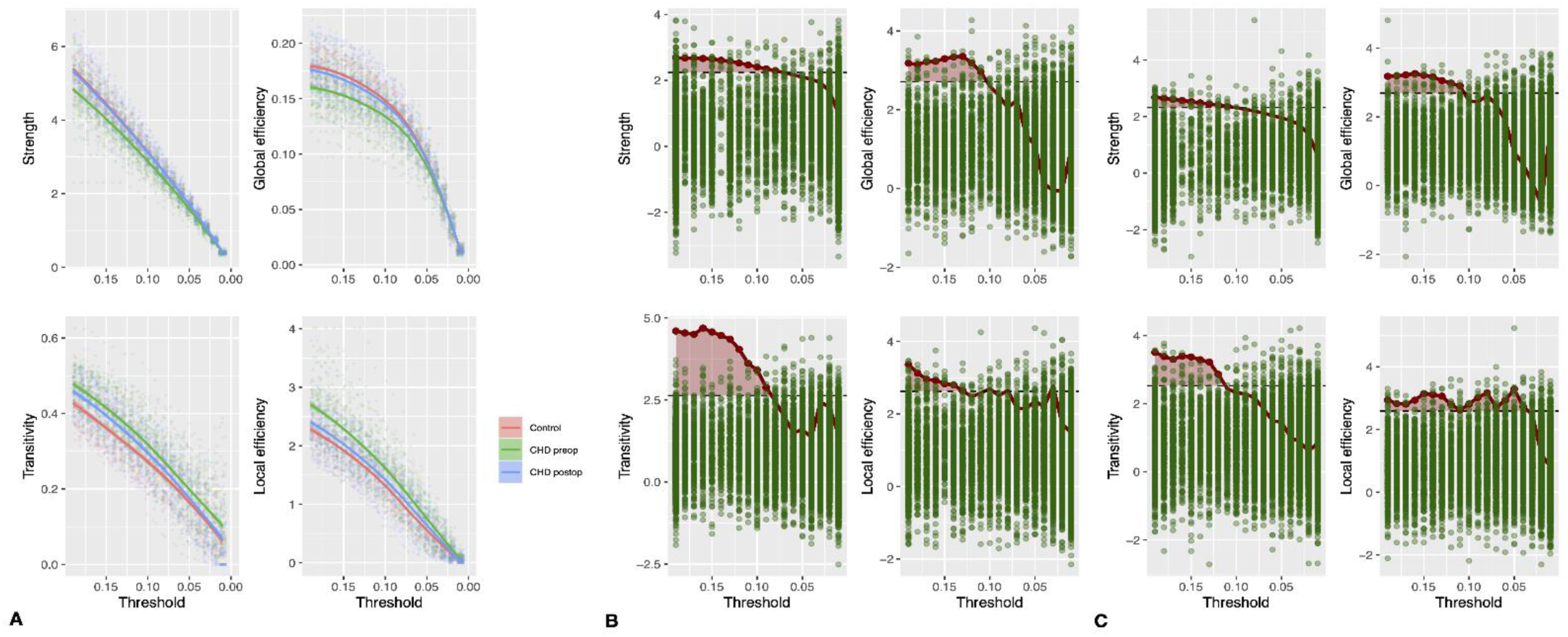
A: Global graph theory parameters in pre- and postoperative CHD neonates and controls across cost threshold range. B Results from MTPC comparison of preoperative CHD neonates and C of postoperative CHD neonates versus healthy controls. Comparison corrected for age at scan, MRI cohort and sex. Area in read indicates threshold range at which group differences were statistically significant (Amtpc), green dots correspond to maximum permuted statistics for each permutation at each threshold (Acrit). Null hypotheses were rejected if Amtpc > Acrit. Here for global efficiency, transitivity and local efficiency. CHD, congenital heart disease; MTPC, multi-threshold permutation correction.

#### Nodal network topology

Preoperatively, Similar differences in nodal level network parameters were found in the comparison of CHD neonates with controls. MTPC revealed higher nodal efficiency and strength in controls compared to preoperative CHD neonates, whereas local efficiency and transitivity were higher in CHD neonates compared to controls (**Fig. 2**). Differences in nodal efficiency and local efficiency involved the frontal, limbic, parietal, occipital and insular lobe and nodes in the subcortical grey matter bilaterally. Higher nodal transitivity in preoperative CHD neonates compared to controls was found predominantly in the limbic lobe in both hemispheres, but affected also parietal, frontal and subcortical regions. Higher strength in controls compared to preoperative CHD neonates was found in multiple nodes in the frontal lobe, in the right precentral gyrus and in the right inferior occipital gyrus. In contrast, significantly higher strength was found in the thalamus bilaterally in preoperative CHD neonates compared to controls. For a complete list of nodal graph theory parameter differences see **Supplementary Table 1**. Postoperatively, when comparing CHD neonates with controls no statistically significant differences in network topology were found at the nodal level.

**Fig. 2.**
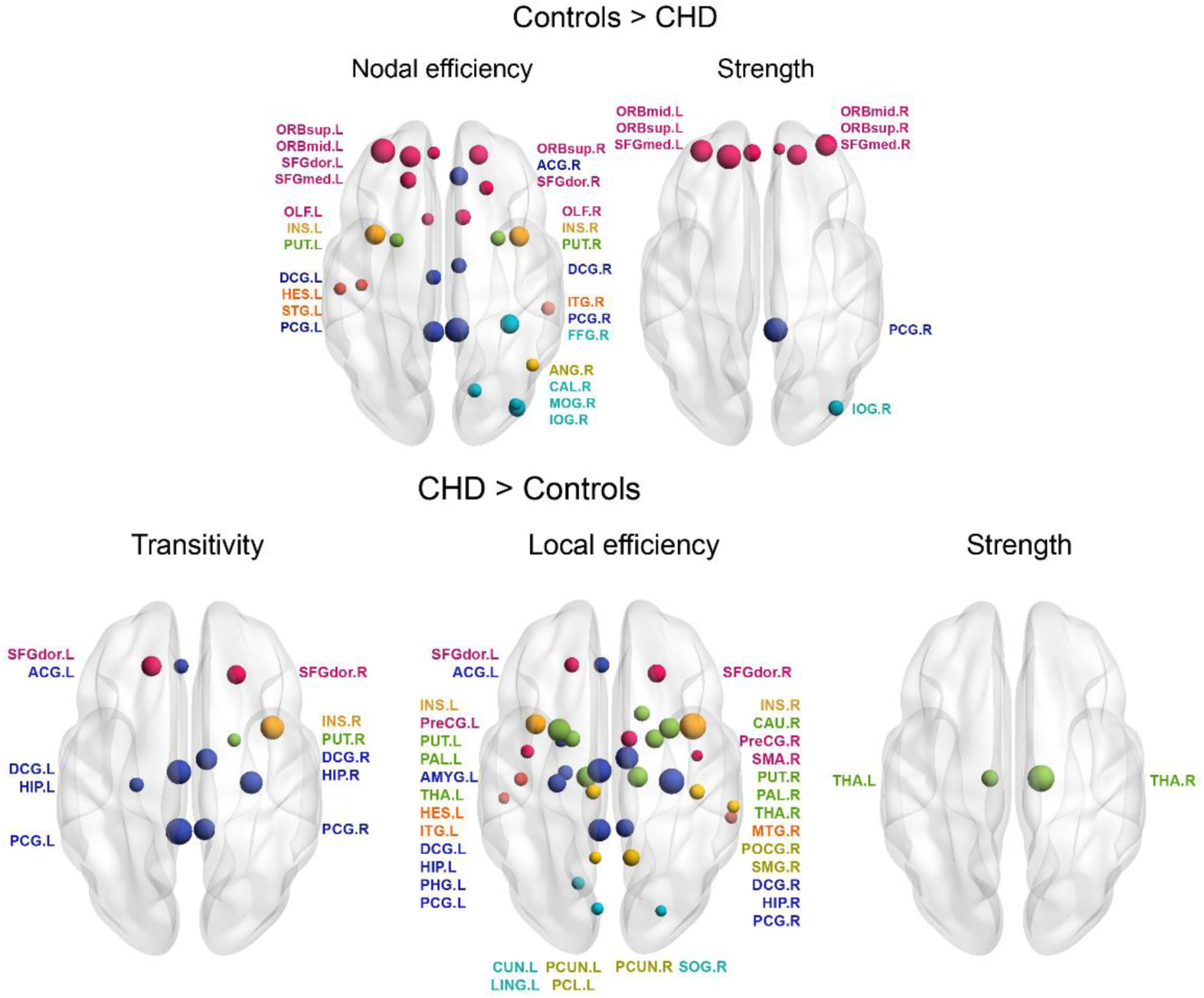
Significant nodal network parameter differences among preoperative CHD neonates and controls tested with MTPC. Comparison corrected for age at scan, MRI cohort and sex. P-values were adjusted for multiple comparison across all 90 anatomical regions of interest with the Benjamini Hochberg procedure. The left hemisphere is displayed on the left side of the image. Nodal size corresponds to the Amtpc value. Nodes are coloured according to lobe membership (pink: frontal; orange: insula; darkblue: limbic; lightblue: occipital; green: subcortical grey matter; yellow: parietal). For a list of node label abbreviations see Supplementary Table 2. CHD, congenital heart disease; MPTC, multi-threshold permutation correction.

#### Edge-level network comparison

NBS carried out at a network cost of 0.19 revealed multiple connected components with significantly different connectivity strength in CHD neonates compared to controls. Preoperatively, two connected subnetworks with 18 nodes sharing 17 edges (P = 0.001), and eight nodes sharing seven edges (P = 0.005) with significantly stronger connections in controls compared to preoperative CHD neonates were found. Of these edges, 66.7% were interhemispheric connections. Furthermore, three connected subnetworks with significantly stronger edges in CHD neonates compared to controls were found (16 nodes, 18 edges, P = 0.001; 11 nodes, 11 edges, p 0.006; 8 nodes, 7 edges, P = 0.009) (**Fig. 3, A-B**). Of these edges 8.33% were interhemispheric connections. Postoperatively, two connected subnetworks were found with significantly higher connectivity strength in controls compared to CHD neonates (9 nodes, 8 edges, P = 0.003; 4 nodes, 3 edges, P = 0.033) (**Fig. 3, C**). Of these edges 54.5% were interhemispheric. Likewise, two subnetworks with stronger connections in CHD compared to controls were found (4 nodes, 3 edges, P = 0.012; 3 nodes, 2 edges, P = 0.04) (**Fig. 3, D**) of which no edges were interhemispheric.

**Figure 3.**
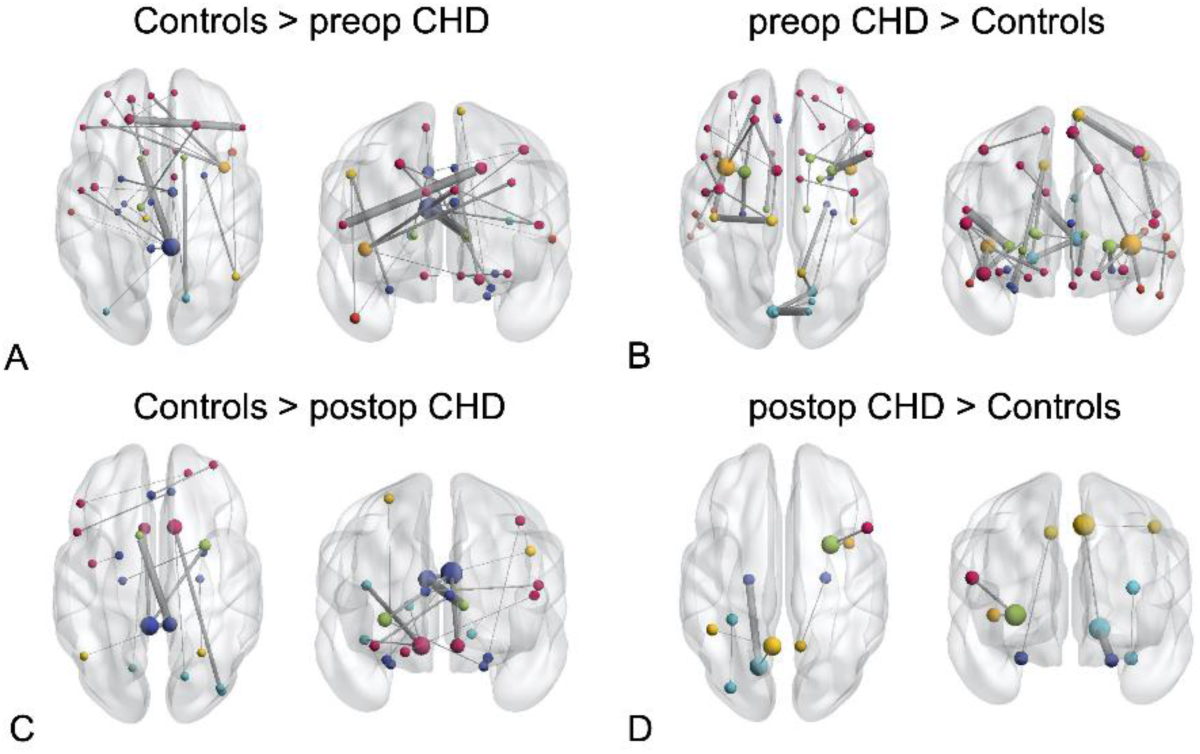
Results of network based statistic showing edgewise network differences between pre-. (A-B) and postoperative CHD (C-D) neonates and healthy controls. A Connected components with higher connectivity strength in controls compared to preoperative CHD neonates. B Components with higher connectivity strength in preoperative CHD neonates compared to controls. C Network components with higher connectivity strength in controls compared to postoperative CHD neonates. D Components with higher connectivity strength in postoperative CHD neonates compared to controls. NBS was carried out at the cost threshold 0.19. Comparison corrected for age at scan, MRI cohort and sex. Axial and coronal views of network components are shown. The left hemisphere is displayed on the left side of the image. Node size corresponds to the nodal degree (number of connections), node colour corresponds to lobe membership (pink: frontal; orange: insula; darkblue: limbic; lightblue: occipital; green: subcortical grey matter; yellow: parietal). Edge size corresponds to test statistic value. CHD, congenital heart disease.

### Trend-level evidence for improvement of structural network topology in class I CHD subgroup

In random coefficient linear mixed effects models the development of network parameters across the first weeks of life was investigated. Global efficiency and strength were positively associated with PMA at scan (ß= 0.015, 95% CI 0.013 to 0.017, P < 0,0001, P_adj_ < 0.0001; ß= 0.44, 95% CI 0.38 to 0.50, P < 0.0001, P_adj_ < 0.0001), whereas the opposite relationship was found for local efficiency and transitivity (ß = −0.26, 95% CI −0.31 to −0.21, P < 0.0001, P_adj_ < 0.0001; ß = −0.018, 95% CI −0.025 to −0.012, P < 0.0001, P_adj_ < 0.0001) (Fig. 4 A). Sex was not significantly associated with network parameters or developmental slopes. An interaction term of PMA at scan and subgroup (levels: control, cardiac class I, class II, class III, class IV) was introduced to test whether the slope of the network parameter development was modified by CHD severity subtype. For local efficiency a significant interaction term was found, suggesting a faster decrease in local efficiency in CHD neonates with class I subtype compared to controls. However, this interaction term did not survive Bonferroni correction for multiple comparison (ß = −0.17, 95% CI −0.32 to −0.013, P = 0.034, P_adj_ = 0.14) (**Fig. 4 B**). Furthermore, there was no difference in slopes among other subclasses compared to controls or compared to each other.

**Figure 4.**
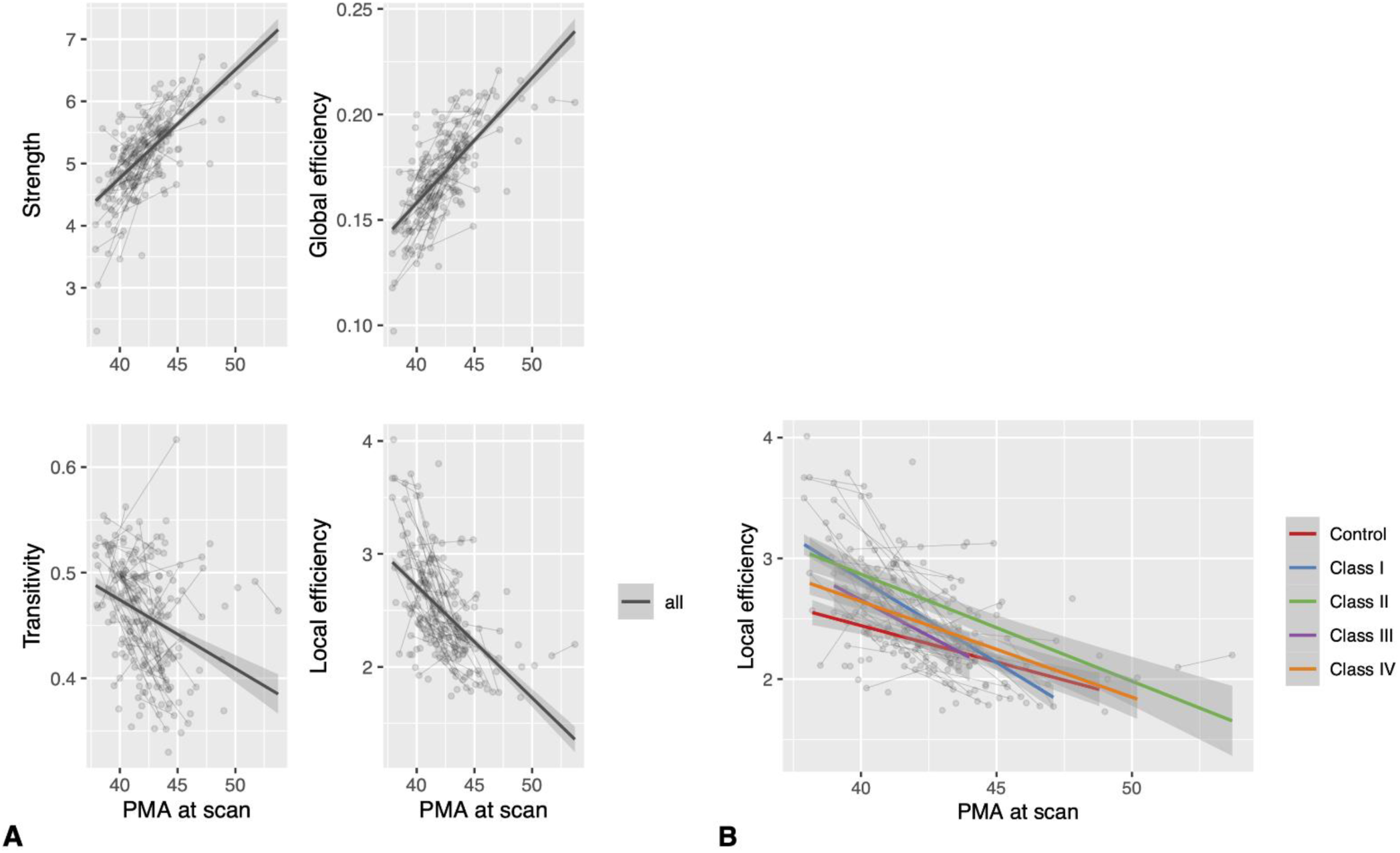
Results from random coefficient mixed models. A Association of global graph theory parameters with PMA at scan including all CHD and control connectomes. B Trajectory of local efficiency development among CHD severity subtypes and healthy controls. Differences in slopes were not statistically significant after correction for multiple comparison. Cardiac subtypes were grouped according to cardiac severity classes by Clancy et al. 2000. PMA, postmenstrual age; CHD, congenital heart disease.

No significant interaction was found for the remaining network parameters. Neither did the grouping of CHD subtypes by arch obstruction or biversus univentricular cardiac physiology show evidence for differences between CHD subtypes.

### WMI is associated with lower strength and integration of global networks across CHD subtypes

In a second random coefficient model including only CHD neonates, an interaction term of cumulative WMI volume and age at scan was introduced to test the effect of WMI burden on network development. Higher cumulative WMI volume was negatively associated with global efficiency (ß = −0.0036, 95% CI - 0.0062 to −0,00092, P = 0.0086, P_adj_ = 0.034) and strength (ß = −0.15, 95% CI −0.24 to −0.055, P = 0.0021, P_adj_ = 0.0084). However, WMI did not modify the effect of age at scan on network parameter development.

### Post-hoc sensitivity analysis

To test the robustness of global graph theory differences between CHD neonates and controls, we repeated the global network comparison after excluding all subjects with WMI or arterial ischemic stroke (n = 18 preoperative, n = 18 postoperative CHD neonates). Preoperatively, global efficiency remained significantly lower, and transitivity remained significantly higher in CHD neonates without brain lesions compared to healthy controls. Postoperatively, no significant differences in global network topology were found between CHD neonates without brain lesions and controls.

## Discussion

In this prospective cohort study, we found less mature structural whole brain connectomes in neonates with CHD compared to healthy controls on all levels of network analysis. The maturational delay was more prominent in the preoperative but persisted to the postoperative period. Trend-level evidence suggested that cardiac physiology might modify the trajectory of perioperative network development. Importantly, larger WMI volume was associated with reduced global efficiency and strength of the whole-brain network topology and was thus identified as a risk factor for perturbation of the neonatal network development.

### Reduced integration and higher segregation of global and nodal network topology in CHD neonates

We found significantly altered structural whole-brain network topology in CHD neonates compared to healthy controls. The differences were evident preoperatively and persisted, albeit less prominent, postoperatively. They can be summarized as a pattern of reduced integration and increased segregation. Similar findings of a perturbated network architecture in CHD neonates compared to controls were reported by Schmithorst et al. (Schmithorst *et al*. 2018). In their study CHD neonates underwent either pre- or postoperative MRI. Pre- and postoperatively reduced global and nodal efficiency, i.e. structural integration were found in FA_mean_ networks, however when controlling for network cost, these differences did not remain significant. In contrast, our proportional cost thresholding paired with MTPC revealed network topological differences between CHD neonates and controls that were independent of network cost. Furthermore, we were able to reveal perturbations in network segregation. In contrast to our analysis, CHD neonates with any type of brain injury were excluded in Schmithorst et al. (Schmithorst *et al*. 2018). We assume that our results are more representative of network disturbances found in CHD neonates as perioperative brain injury, such as punctate WMI and focal stroke, are common in CHD (Peyvandi *et al*. 2019). However, the different findings in our study and the analysis by Schmithorst et al. cannot be entirely explained by the cases with brain injury as revealed by our post-hoc sensitivity analysis. We found that after exclusion of cases with injury preoperative global network differences remained, while postoperatively significant differences did not. Beside the effects of injury, the loss of postoperative differences might have also resulted from reduced sample size and statistical power.

Opposed to our structural connectomics findings, a previous analysis of the functional connectome demonstrated a preserved functional global network architecture in CHD neonates (De Asis-Cruz *et al*. 2018). This contrast in global topological findings could be partly attributed to the exclusion of CHD neonates with parenchymal brain lesions, but could as well arise from temporal differences in functional and structural network maturation. At the neonatal age structural precedes functional network development, whereas the coupling of both networks gradually increases from the last trimester of gestation to 20 years of age (Cao *et al*. 2017). Therefore, structural might precede functional connectomic disturbances, with the latter only becoming apparent at later ages.

Our network edge-level analysis revealed multiple subnetworks that differed between CHD neonates and controls pre- and postoperatively, with interhemispheric connections appearing most vulnerable to perturbations in CHD. Impaired interhemispheric connectivity evident as volumetric and microstructural dysmaturation in the corpus callosum have been reported in a small subsample of our CHD cohort (Hagmann *et al*. 2016). Our findings also accord with a study showing that approximately 60% of functional connections that were weaker in CHD neonates than in controls were interhemispheric (De Asis-Cruz *et al*. 2018).

### Connectome topology reflects structural network dysmaturation in CHD neonates

The longitudinal trajectory of network development in our cohort reflects that global efficiency and strength increased significantly across the investigated period of neonatal development, whereas parameters of segregation such as local efficiency and transitivity significantly decreased. This is in line with the suggested normative trajectory of network development during foetal and neonatal brain development, characterized by an increase in network integration (i.e. global efficiency) and decrease in network segregation (i.e. local efficiency, transitivity) (Takahashi *et al*. 2012, Tymofiyeva *et al*. 2013, Jakab *et al*. 2014, Song *et al*. 2017, Keunen *et al*. 2018, Turk *et al*. 2019). Axonal growth, synaptogenesis and myelination starting during the late second and early third trimester of gestation lead to the emergence of interhemispheric connections and intrahemispheric long-range association fibres (Keunen *et al*. 2017) and contribute to an increasing structural integration of the whole brain network (Tymofiyeva *et al*. 2013, Song *et al*. 2017). This increasing efficiency is paralleled by decreasing segregation as result of synaptic pruning and the refinement of the specialized subnetworks (Damaraju *et al*. 2014). Given this pattern of structural connectome development, our finding of reduced global efficiency, and higher local efficiency and transitivity in CHD neonates indicates a maturational delay in network architecture. These observations in the early preoperative period suggest that network dysmaturation in CHD neonates might be of foetal origin with a persistence into the postoperative period. Furthermore, our findings demonstrate an impairment beyond the previously described multi-faceted brain developmental delay in CHD neonates that involves structural and metabolic aberrations and originates in the last trimester of gestation (Limperopoulos *et al*. 2010, Berman *et al*. 2011, Clouchoux *et al*. 2013).

### Cardiac physiology might modify perioperative network development in CHD neonates

The trajectories of brain development in CHD neonates can be modified by the type of cardiac defect and postoperative cardiac physiology as shown previously (Peyvandi *et al*. 2018). When comparing trajectories between CHD severity subgroups and controls, we found trend-level evidence for a difference in local efficiency development between class I CHD neonates and controls. More rapid decrease of local efficiency from the pre- to the postoperative period could suggest a catch up in network development in this CHD subclass on the mild end of the severity spectrum. The majority of class I CHD neonates had a transposition of the great arteries, and thus received surgical restoration of normal cardiac physiology. This improvement in network topology from the pre- to the postoperative period evident in the largest CHD subgroup could drive our observation that the difference in network topology between CHD neonates and controls was less prominent postoperatively. However, when conservatively correcting for multiple comparison, this difference did not remain significant. While this is consistent with similar white matter micro-structural change rates in neonates with different subtypes of CHD (Claessens *et al*. 2019), further research with even larger and more balanced subgroups is needed to explore the modifying effect of cardiac physiology on network development.

### WMI is associated with widespread network perturbations

We found evidence for a negative effect of WMI burden on global efficiency and strength of the structural whole brain network in CHD neonates, identifying WMI as potential risk factor for network perturbations. This disruptive effect of WMI on connectivity is in line with functional connectomics studies in preterm born infants, revealing a negative association between WMI and thalamocortical (Duerden *et al*. 2019) as well as interhemispheric connectivity (Smyser *et al*. 2013).

Given the differences in spatial distribution of WMI in CHD compared to preterm infants, with a paucity of central WMI in CHD (Guo *et al*. 2019), it seems striking that the mainly small punctate lesions have such widespread effects on global network integration and strength in CHD neonates. Whereas the impact of lesion location on network dysconnectivity warrants further investigation, this could highlight that the overt WMI depicted on diagnostic MRI is accompanied by covert and widespread microstructural white matter dysmaturation (Dimitropoulos *et al*. 2013). The latter is only evident on advanced neuroimaging and could explain the systems-level effects of WMI observed in our study. There is evidence that WMI furthermore impacts longitudinal micro- and macrostructural brain development in CHD infants with moderate to severe WMI or stroke when compared to infants without ischemic injury (Claessens *et al*. 2019). Moreover, brain growth was found to be impaired in CHD neonates with moderate to severe WMI (Peyvandi *et al*. 2018). In our cohort, covering a narrow range of age at scan, we did not find a modifying effect of WMI on the slope of network development.

### Similar findings in other age groups and potential functional relevance

In line with our findings, a previous study found trend-level reduced global efficiency and significantly higher modularity, indicative of reduced structural integration and increased segregation, in *adolescents* with transposition of the great arteries (Panigrahy *et al*. 2015). Together our findings suggest that neonatal network perturbations might persist into adolescence, however this has yet to be tested in a longitudinal dataset. Furthermore, Panigrahy et al. demonstrated that disrupted network organization in CHD *adolescent* is of functional relevance as network topology was found to mediate cognitive performance across multiple domains (Panigrahy *et al*. 2015). However, how the altered neonatal network architecture in CHD infants impacts later neurodevelopment remains to be elucidated.

### Limitations

The following limitations of our study need to be mentioned. Our analysis is based on DTI tractography, which is inevitably an imperfect reconstruction of complex WMI fibre tract architecture (Tymofiyeva *et al*. 2012). The neonatal brain is still undergoing myelination and many of the white matter structures underlying cortico-cortical connectivity are unmyelinated, which leads to lower diffusion anisotropy than in children or adults. Lower anisotropy results in higher uncertainty when estimating fibre orientations, which is particularly challenging in crossing-fibre regions. While our study protocol represents a realistic compromise between image quality and scan time in this sensitive patient population, studies using higher angular resolution datasets will likely provide better estimates of whole-brain structural connectivity.

As only two imaging time points were acquired in our cohort the exploration and modelling of more complex growth trajectories was hindered. However, other aspects of brain development such as structural brain growth and functional network development do not follow linear but rather asymptotic growth trajectories (Dosenbach *et al*. 2010, Holland *et al*. 2014). This might be particularly true during the first months of life, as structural brain growth shows a tapering off of the daily growth rate at about 3 months (Holland *et al*. 2014). Thus, we expect that the development of network parameters during this time might follow more complex growth trajectories too. More densely sampled longitudinal neuroimaging data across a longer timespan would be necessary to establish and investigate more complex growth trajectories of structural network architecture in healthy and aberrant neurodevelopment. However, in a typical clinical setting, this remains a challenging goal particularly for neonates with critical CHD.

In a small fraction of included CHD neonates in our study arterial ischemic strokes were observed. These strokes were mainly focal and restricted to the cortex. As the group of neonates with stroke was very small, no further testing of the effect of stroke on the structural connectome was performed. This remains to be addressed in future research.

A further limitation of the interpretation of our findings is that results might depend on the connectomics approach deployed. For instance, the parcellation scheme, the tractography algorithm, or the way cost-thresholding is applied. It remains a challenge of this rapidly emerging field to establish methodological standards and ensure replicability and comparability of results (Tymofiyeva *et al*. 2012). Further reproducibility studies are needed to help depict and control technical bias.

## Conclusion

In our cohort of neonates with CHD undergoing cardiopulmonary bypass surgery we found evidence for a maturational delay in the structural brain connectome characterized by lower integration and higher segregation. This network dysmaturation was most prominent in the preoperative but persisted to the postoperative period. We found trend-level evidence for a modifying effect of CHD severity subtype on network development, suggesting that postoperative normalization of cardiac physiology may improve structural network topology. WMI burden was associated with perturbation of the structural connectome, impacting global network strength and efficiency. Our findings advance our understanding of aberrant brain development in CHD neonates. We revealed potential risk factors for aberrant network development, and mechanisms how these findings could represent neural correlates of later neurodevelopmental sequelae.

## Acknowledgements

The authors would like to thank the patients and families who participated in this study. Furthermore, we would like to thank Dr. Shabnam Peyvandi from the UCSF Benioff Children’s Hospital for her advice on building mixed models for longitudinal neuroimaging analyses.

**MF** was supported by the Young Investigator Exchange Programme of the European Society for Paediatric Research and the Anna Müller Grocholski Foundation. **AJ** was supported by the FZK Foundation, Swiss National Science Foundation SPARK Grant, the OPO Foundation, the Anna Müller Grocholski Foundation and the Prof. Dr. Max Cloetta Foundation. **SPM** is supported by the Bloorview Children’s Hospital Chair in Paediatric Neuroscience.

**This manuscript is the author’s original version**.

## Supplementary Materials

### Graph theory parameters

#### Global and nodal efficiency

The networks ability to transfer information in parallel was quantified by calculating the weighted global efficiency, which is the reciprocal of the harmonic mean of the shortest weighted path length. Networks that have a short average shortest path length between two nodes are considered topologically integrated and efficient (Latora et al. 2001). Global efficiency for a weighted graph *G* was calculated based on the following equation

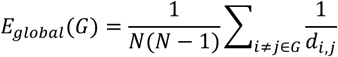

where *N* is the total number of nodes in the network *G*;*d*_*i,j*_ is the weighted shortest paths length between nodes *i* and *j* (Latora et al. 2003). Accordingly, weighted nodal efficiency was calculated as nodal counterpart of the weighted global efficiency. Nodal efficiency quantifies the integration of a specific node with all other nodes in the network. It is the normalized sum of the inverse of the shortest path length *d*_*i,j*_ of a specific node *i* to all other nodes (Latora et al. 2003).

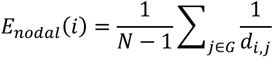

We expect global and nodal efficiency to increase during neonatal brain development (Cao et al. 2017).

#### Local efficiency

Local efficiency as opposed to nodal efficiency quantifies the integration of a specific node within a subgraph *G*_*i*_, encompassing all nodes that are immediate neighbours of the node (Latora *et al*. 2001, Latora *et al*. 2003). Thus, local efficiency is a measure of structural segregation.

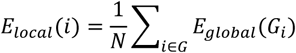

The average local efficiency across all nodes serves as global measure of the local efficiency of a network. Local efficiency is expected to decrease during neonatal brain development (Cao *et al*. 2017).

#### Strength

By summing the edge weights (*w*_*i,j*_) of all edges connected to a node, nodal strength was calculated. Global strength was measured by averaging the strength of all nodes in the network (Barrat et al. 2004) and is expected to increase with brain network development.

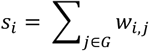

#### Transitivity

Transitivity, which is sometimes also referred to as clustering coefficient, quantifies the probability that the nodes *j* and *h*, which are directly connected to node *i* are also directly connected with each other (Barrat et al. 2004). For global transitivity the number of connected triangles is divided by the number of connected triplets. For the nodal counterpart the weighted transitivity was calculated following the generalization by Barrat et al. (Barrat et al. 2004). *w*_*i,h*_ is the edge weight; *s*_*i*_ is the sum of all neighboring edge weights; *k*_*i*_ is the vertex degree, i.e. the number of all edges connected to that node; *a*_*ij*_ denotes the element in the adjacency matrix.

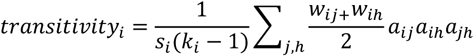

Transitivity as measure of structural segregation is expected to decrease with brain network development (Cao et al. 2017).

#### Multi-threshold permutation correction

To test group differences with Multi-threshold permutation correction (MTPC), general linear models (GLM) were calculated at each individual cost threshold and test statistics were noted. By permuting the group assignment in the GLM across n iterations the test-statistics under the null hypothesis were sampled for each threshold and the maximum of the null test-statistics for each permutation was determined. The critical test-statistics threshold above which the results were significant at the specified alpha-level were identified. For each cluster of ≥ 3 consecutive thresholds with test-statistics above the critical test-statistics threshold the area under the curve (AUC), denoted as *AUC*_*mtpc*_, was calculated. Similarly, the mean of the super-critical AUCs of the permuted statistics under the null hypothesis was calculated (*AUC*_*crit*_). The null-hypothesis was rejected if *AUC*_*mtpc*_>*AUC*_*crit*_. The number of permutations n was set to 5000 for nodal and 10 000 for global graph metric comparisons for each threshold as suggested by Watson et al. (Watson et al. 2019).

---

**Supplementary Figure 1.**
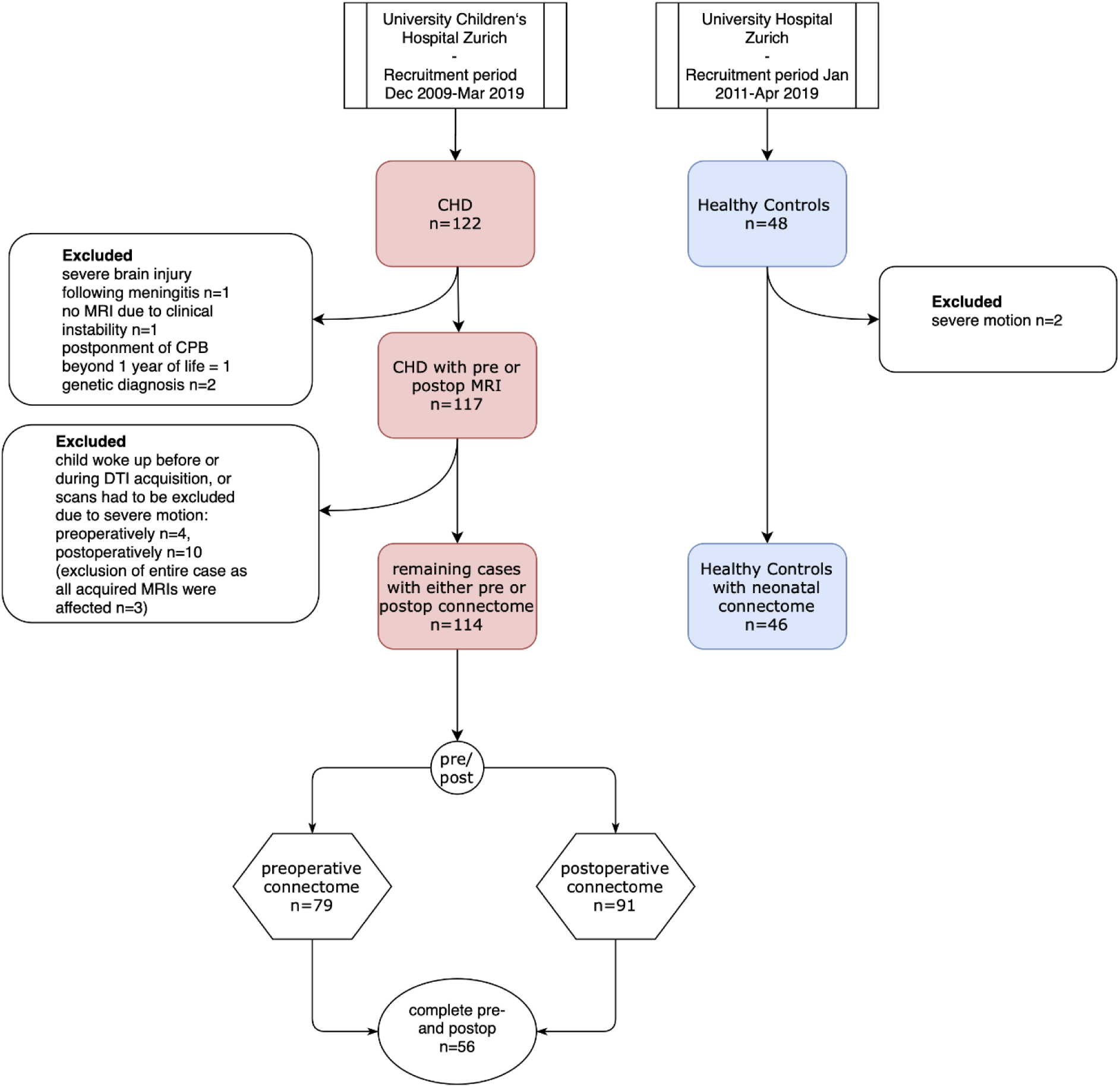
Flow chart of included subjects and available connectomes.

**Supplementary Table 1.**
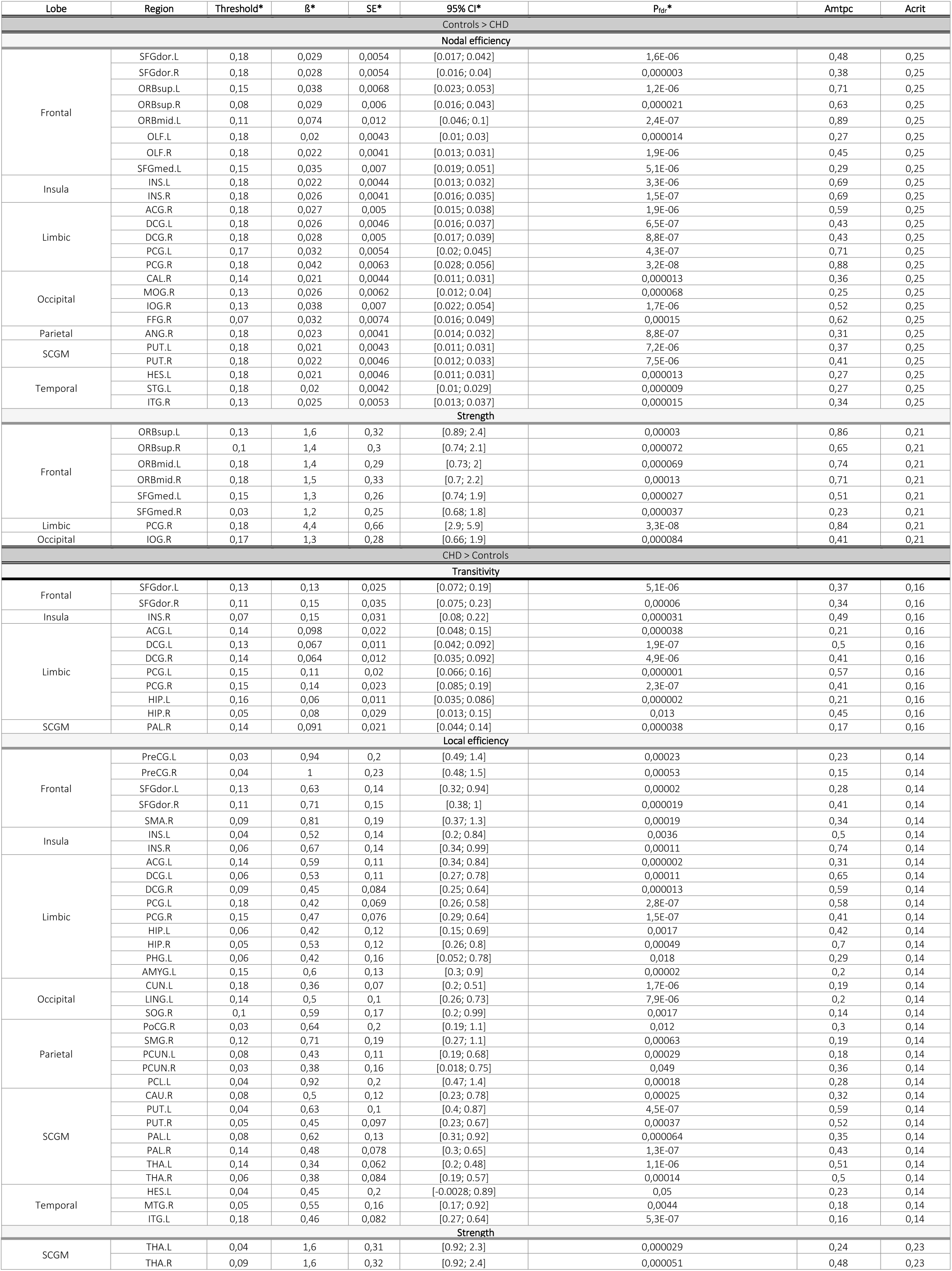
Significant nodal level differences between preoperative CHD neonates and healthy controls as revealed by MTPC. As two one-sided tests were performed to determine the direction of the effects, results are grouped by the contrast “Controls > CHD” or “CHD > Controls”. *Threshold indicates the cost threshold at which the strongest *ß coefficient was observed. Statistical parameters are given for that threshold. *P_fdr_ is the threshold-specific P-value corrected for multiple comparison across all 90 ROIs by means of the Benjamini Hochberg procedure. Amtpc and Acrit denote the results of the overall MTPC comparison across the whole range of thresholds. SCGM, subcortical gray matter.

**Supplementary Table 2.**
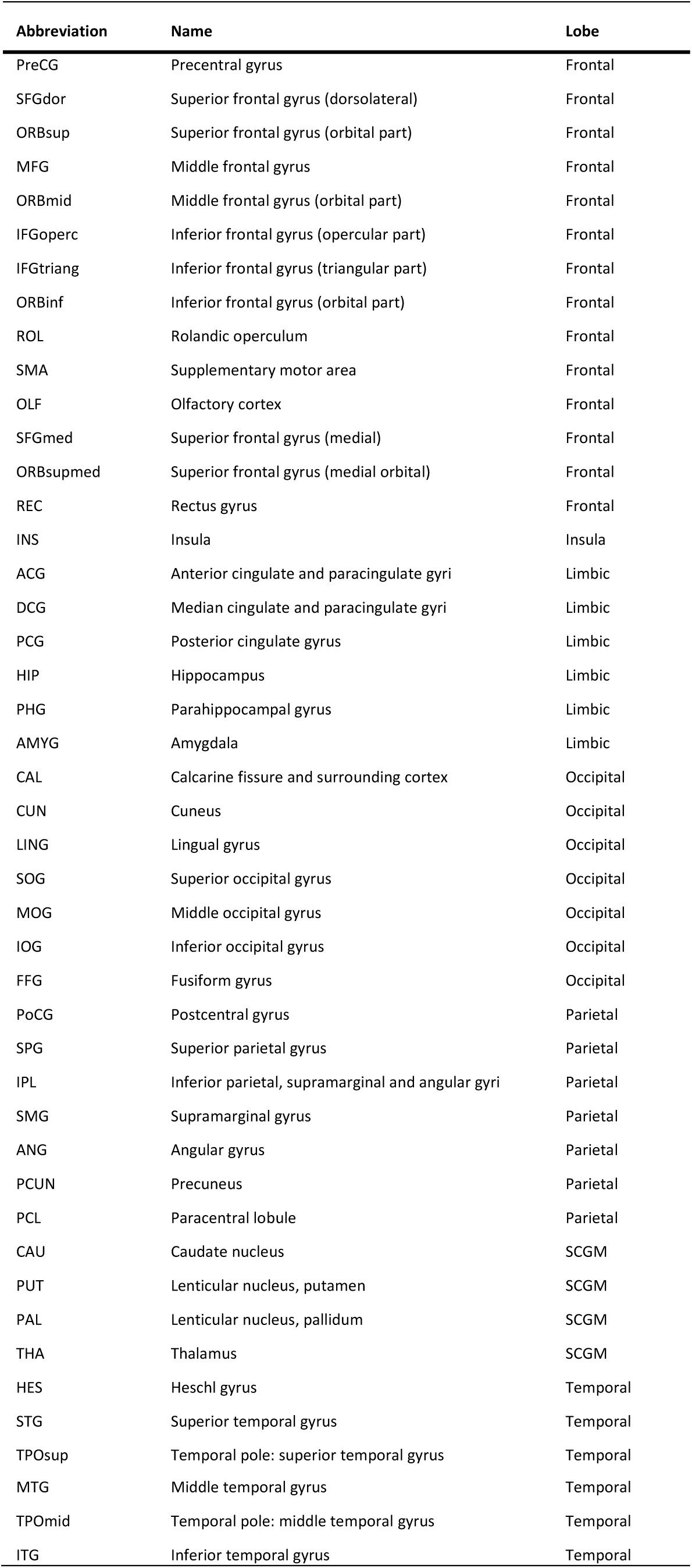
List of node abbreviations parcellated with the Automated Anatomical Labeling. Abbreviations are appended by .L for left hemispheric and .R for right hemispheric regions.

## References

Andropoulos, D. B., H. B. Ahmad, T. Haq, K. Brady, S. A. Stayer, M. R. Meador, J. V. Hunter, C. Rivera, R. G. Voigt, M. Turcich, C. Q. He, L. S. Shekerdemian, H. A. Dickerson, C. D. Fraser, E. Dean McKenzie, J. S. Heinle and R. Blaine Easley (2014). “The association between brain injury, perioperative anesthetic exposure, and 12-month neurodevelopmental outcomes after neonatal cardiac surgery: a retrospective cohort study.” Paediatr Anaesth 24(3): 266–274.

Batalle, D., E. J. Hughes, H. Zhang, J. D. Tournier, N. Tusor, P. Aljabar, L. Wali, D. C. Alexander, J. V. Hajnal, C. Nosarti, A. D. Edwards and S. J. Counsell (2017). “Early development of structural networks and the impact of prematurity on brain connectivity.” Neuroimage 149: 379–392.

Benjamini, Y. and Y. Hochberg (1995). “Controlling the False Discovery Rate - a Practical and Powerful Approach to Multiple Testing.” Journal of the Royal Statistical Society Series B-Statistical Methodology 57(1): 289–300.

Berman, J. I., S. E. Hamrick, P. S. McQuillen, C. Studholme, D. Xu, R. G. Henry, L. K. Hornberger and O. A. Glenn (2011). “Diffusion-weighted imaging in fetuses with severe congenital heart defects.” AJNR Am J Neuroradiol 32(2): E21–22.

Bertholdt, S., B. Latal, R. Liamlahi, R. Pretre, I. Scheer, R. Goetti, H. Dave, V. Bernet, A. Schmitz, M. von Rhein, W. Knirsch, H. Research Group and Brain (2014). “Cerebral lesions on magnetic resonance imaging correlate with preoperative neurological status in neonates undergoing cardiopulmonary bypass surgery.” Eur J Cardiothorac Surg 45(4): 625–632.

Cai, Y., X. Wu, Z. Su, Y. Shi and J.-H. Gao (2017). “Functional thalamocortical connectivity development and alterations in preterm infants during the neonatal period.” Neuroscience.

Cao, M., H. Huang and Y. He (2017). “Developmental Connectomics from Infancy through Early Childhood.” Trends Neurosci 40(8): 494–506.

Ceschin, R., V. K. Lee, V. Schmithorst and A. Panigrahy (2015). “Regional vulnerability of longitudinal cortical association connectivity: Associated with structural network topology alterations in preterm children with cerebral palsy.” Neuroimage Clin 9: 322–337.

Claessens, N. H., P. Moeskops, A. Buchmann, B. Latal, W. Knirsch, I. Scheer, I. Isgum, L. S. de Vries, M. J. Benders and M. von Rhein (2016). “Delayed cortical gray matter development in neonates with severe congenital heart disease.” Pediatr Res 80(5): 668–674.

Claessens, N. H. P., S. O. Algra, T. L. Ouwehand, N. J. G. Jansen, R. Schappin, F. Haas, M. J. C. Eijsermans, L. S. de Vries, M. Benders and C. H. D. L. S. G. Utrecht (2018). “Perioperative neonatal brain injury is associated with worse school-age neurodevelopment in children with critical congenital heart disease.” Dev Med Child Neurol 60(10): 1052–1058.

Claessens, N. H. P., J. Breur, F. Groenendaal, R. M. Wosten-van Asperen, R. Stegeman, F. Haas, J. Dudink, L. S. de Vries, N. J. G. Jansen and M. Benders (2019). “Brain microstructural development in neonates with critical congenital heart disease: An atlas-based diffusion tensor imaging study.” Neuroimage Clin 21: 101672.

Claessens, N. H. P., C. J. Kelly, S. J. Counsell and M. Benders (2017). “Neuroimaging, cardiovascular physiology, and functional outcomes in infants with congenital heart disease.” Dev Med Child Neurol 59(9): 894–902.

Clancy, R. R., S. A. McGaurn, G. Wernovsky, T. L. Spray, W. I. Norwood, M. L. Jacobs, J. D. Murphy, J. W. Gaynor and J. E. Goin (2000). “Preoperative risk-of-death prediction model in heart surgery with deep hypothermic circulatory arrest in the neonate.” J Thorac Cardiovasc Surg 119(2): 347–357.

Clouchoux, C., A. J. du Plessis, M. Bouyssi-Kobar, W. Tworetzky, D. B. McElhinney, D. W. Brown, A. Gholipour, D. Kudelski, S. K. Warfield, R. J. McCarter, R. L. Robertson, Jr., A. C. Evans, J. W. Newburger and C. Limperopoulos (2013). “Delayed cortical development in fetuses with complex congenital heart disease.” Cereb Cortex 23(12): 2932–2943.

Csardi, G. and T. Nepusz (2006). “The igraph software package for complex network research.” InterJournal Complex Systems: 1695.

Damaraju, E., A. Caprihan, J. R. Lowe, E. A. Allen, V. D. Calhoun and J. P. Phillips (2014). “Functional connectivity in the developing brain: a longitudinal study from 4 to 9months of age.” Neuroimage 84: 169–180.

De Asis-Cruz, J., M. T. Donofrio, G. Vezina and C. Limperopoulos (2018). “Aberrant brain functional connectivity in newborns with congenital heart disease before cardiac surgery.” Neuroimage Clin 17: 31–42.

Dimitropoulos, A., P. S. McQuillen, V. Sethi, A. Moosa, V. Chau, D. Xu, R. Brant, A. Azakie, A. Campbell, A. J. Barkovich, K. J. Poskitt and S. P. Miller (2013). “Brain injury and development in newborns with critical congenital heart disease.” Neurology 81(3): 241–248.

Dosenbach, N. U., B. Nardos, A. L. Cohen, D. A. Fair, J. D. Power, J. A. Church, S. M. Nelson, G. S. Wig, A. C. Vogel, C. N. Lessov-Schlaggar, K. A. Barnes, J. W. Dubis, E. Feczko, R. S. Coalson, J. R. Pruett, Jr., D. M. Barch, S. E. Petersen and B. L. Schlaggar (2010). “Prediction of individual brain maturity using fMRI.” Science 329(5997): 1358–1361.

Drakesmith, M., K. Caeyenberghs, A. Dutt, G. Lewis, A. S. David and D. K. Jones (2015). “Overcoming the effects of false positives and threshold bias in graph theoretical analyses of neuroimaging data.” Neuroimage 118: 313–333.

Duerden, E. G., S. Halani, K. Ng, T. Guo, J. Foong, T. J. A. Glass, V. Chau, H. M. Branson, J. G. Sled, H. E. Whyte, E. N. Kelly and S. P. Miller (2019). “White matter injury predicts disrupted functional connectivity and microstructure in very preterm born neonates.” Neuroimage Clin 21: 101596.

Ferry, P. C. (1987). “Neurologic sequelae of cardiac surgery in children.” Am J Dis Child 141(3): 309–312.

Guo, T., V. Chau, S. Peyvandi, B. Latal, P. S. McQuillen, W. Knirsch, A. Synnes, M. Feldmann, N. Naef, M. M. Chakravarty, A. De Petrillo, E. G. Duerden, A. J. Barkovich and S. P. Miller (2019). “White matter injury in term neonates with congenital heart diseases: Topology & comparison with preterm new-borns.” Neuroimage 185: 742–749.

Hagmann, C., J. Singer, B. Latal, W. Knirsch and M. Makki (2016). “Regional Microstructural and Volumetric Magnetic Resonance Imaging (MRI) Abnormalities in the Corpus Callosum of Neonates With Congenital Heart Defect Undergoing Cardiac Surgery.” J Child Neurol 31(3): 300–308.

Holland, D., L. Chang, T. M. Ernst, M. Curran, S. D. Buchthal, D. Alicata, J. Skranes, H. Johansen, A. Hernandez, R. Yamakawa, J. M. Kuperman and A. M. Dale (2014). “Structural growth trajectories and rates of change in the first 3 months of infant brain development.” JAMA Neurol 71(10): 1266–1274.

Huisenga, D., S. La Bastide-Van Gemert, A. Van Bergen, J. Sweeney and M. Hadders-Algra (2020). “Developmental outcomes after early surgery for complex congenital heart disease: a systematic review and meta-analysis.” Dev Med Child Neurol.

Jakab, A. (2019). “Developmental Pathoconnectomics and Advanced Fetal MRI.” Top Magn Reson Imaging 28(5): 275–284.

Jakab, A., E. Meuwly, M. Feldmann, M. V. Rhein, R. Kottke, R. O’Gorman Tuura, B. Latal, W. Knirsch, H. Research Group and Brain (2019). “Left temporal plane growth predicts language development in newborns with congenital heart disease.” Brain 142(5): 1270–1281.

Jakab, A., C. Ruegger, H. U. Bucher, M. Makki, P. S. Huppi, R. Tuura, C. Hagmann and E. P. O. N. T. G. Swiss (2019). “Network based statistics reveals trophic and neuroprotective effect of early high dose erythropoetin on brain connectivity in very preterm infants.” Neuroimage Clin 22: 101806.

Jakab, A., E. Schwartz, G. Kasprian, G. M. Gruber, D. Prayer, V. Schopf and G. Langs (2014). “Fetal functional imaging portrays heterogeneous development of emerging human brain networks.” Front Hum Neurosci 8: 852.

Karsdorp, P. A., W. Everaerd, M. Kindt and B. J. Mulder (2007). “Psychological and cognitive functioning in children and adolescents with congenital heart disease: a meta-analysis.” J Pediatr Psychol 32(5): 527–541.

Keunen, K., S. J. Counsell and M. Benders (2017). “The emergence of functional architecture during early brain development.” Neuroimage 160: 2–14.

Keunen, K., H. K. van der Burgh, M. A. de Reus, P. Moeskops, R. Schmidt, L. J. Stolwijk, S. C. de Lange, I. Isgum, L. S. de Vries, M. J. Benders and M. P. van den Heuvel (2018). “Early human brain development: insights into macroscale connectome wiring.” Pediatr Res 84(6): 829–836.

Licht, D. J., D. M. Shera, R. R. Clancy, G. Wernovsky, L. M. Montenegro, S. C. Nicolson, R. A. Zimmerman, T. L. Spray, J. W. Gaynor and A. Vossough (2009). “Brain maturation is delayed in infants with complex congenital heart defects.” J Thorac Cardiovasc Surg 137(3): 529-536; discussion 536-527.

Limperopoulos, C., W. Tworetzky, D. B. McElhinney, J. W. Newburger, D. W. Brown, R. L. Robertson, Jr., N. Guizard, E. McGrath, J. Geva, D. Annese, C. Dunbar-Masterson, B. Trainor, P. C. Laussen and A. J. du Plessis (2010). “Brain volume and metabolism in fetuses with congenital heart disease: evaluation with quantitative magnetic resonance imaging and spectroscopy.” Circulation 121(1): 26–33.

Meuwly, E., M. Feldmann, W. Knirsch, M. von Rhein, K. Payette, H. Dave, R. O. G. Tuura, R. Kottke, C. Hagmann, B. Latal, A. Jakab, H. Research Group and Brain* (2019). “Postoperative brain volumes are associated with one-year neurodevelopmental outcome in children with severe congenital heart disease.” Sci Rep 9(1): 10885.

Miller, S. P., P. S. McQuillen, S. Hamrick, D. Xu, D. V. Glidden, N. Charlton, T. Karl, A. Azakie, D. M. Ferriero, A. J. Barkovich and D. B. Vigneron (2007). “Abnormal brain development in newborns with congenital heart disease.” N Engl J Med 357(19): 1928–1938.

Mulkey, S. B., X. Ou, R. H. Ramakrishnaiah, C. M. Glasier, C. J. Swearingen, M. S. Melguizo, V. L. Yap, M. L. Schmitz and A. T. Bhutta (2014). “White matter injury in newborns with congenital heart disease: a diffusion tensor imaging study.” Pediatr Neurol 51(3): 377–383.

Ortinau, C., J. Beca, J. Lambeth, B. Ferdman, D. Alexopoulos, J. S. Shimony, M. Wallendorf, J. Neil and T. Inder (2012). “Regional alterations in cerebral growth exist preoperatively in infants with congenital heart disease.” J Thorac Cardiovasc Surg 143(6): 1264–1270.

Pandit, A. S., E. Robinson, P. Aljabar, G. Ball, I. S. Gousias, Z. Wang, J. V. Hajnal, D. Rueckert, S. J. Counsell, G. Montana and A. D. Edwards (2014). “Whole-brain mapping of structural connectivity in infants reveals altered connection strength associated with growth and preterm birth.” Cereb Cortex 24(9): 2324–2333.

Panigrahy, A., V. J. Schmithorst, J. L. Wisnowski, C. G. Watson, D. C. Bellinger, J. W. Newburger and M. J. Rivkin (2015). “Relationship of white matter network topology and cognitive outcome in adolescents with d-transposition of the great arteries.” Neuroimage Clin 7: 438–448.

Parker, G. J., H. A. Haroon and C. A. Wheeler-Kingshott (2003). “A framework for a streamline-based probabilistic index of connectivity (PICo) using a structural interpretation of MRI diffusion measurements.” J Magn Reson Imaging 18(2): 242–254.

Partridge, S. C., D. B. Vigneron, N. N. Charlton, J. I. Berman, R. G. Henry, P. Mukherjee, P. S. McQuillen, T. R. Karl, A. J. Barkovich and S. P. Miller (2006). “Pyramidal tract maturation after brain injury in newborns with heart disease.” Ann Neurol 59(4): 640–651.

Peyvandi, S., V. De Santiago, E. Chakkarapani, V. Chau, A. Campbell, K. J. Poskitt, D. Xu, A. J. Barkovich, S. Miller and P. McQuillen (2016). “Association of Prenatal Diagnosis of Critical Congenital Heart Disease With Postnatal Brain Development and the Risk of Brain Injury.” JAMA Pediatr 170(4): e154450.

Peyvandi, S., H. Kim, J. Lau, A. J. Barkovich, A. Campbell, S. Miller, D. Xu and P. McQuillen (2018). “The association between cardiac physiology, acquired brain injury, and postnatal brain growth in critical congenital heart disease.” J Thorac Cardiovasc Surg 155(1): 291–300 e293.

Peyvandi, S., B. Latal, S. P. Miller and P. S. McQuillen (2019). “The neonatal brain in critical congenital heart disease: Insights and future directions.” Neuroimage 185: 776–782.

R Core Team (2019). R: A Language and Environment for Statistical Computing. Vienna, Austria, R Foundation for Statistical Computing.

Rubinov, M. and O. Sporns (2010). “Complex network measures of brain connectivity: uses and interpretations.” Neuroimage 52(3): 1059–1069.

Schielzeth, H. and W. Forstmeier (2009). “Conclusions beyond support: overconfident estimates in mixed models.” Behav Ecol 20(2): 416–420.

Schmithorst, V. J., J. K. Votava-Smith, N. Tran, R. Kim, V. Lee, R. Ceschin, H. Lai, J. A. Johnson, J. S. De Toledo, S. Bluml, L. Paquette and A. Panigrahy (2018). “Structural network topology correlates of microstructural brain dysmaturation in term infants with congenital heart disease.” Hum Brain Mapp 39(11): 4593–4610.

Shi, F., P. T. Yap, G. Wu, H. Jia, J. H. Gilmore, W. Lin and D. Shen (2011). “Infant brain atlases from neonates to 1- and 2-year-olds.” PLoS One 6(4): e18746.

Smyser, C. D., A. Z. Snyder, J. S. Shimony, T. M. Blazey, T. E. Inder and J. J. Neil (2013). “Effects of white matter injury on resting state fMRI measures in prematurely born infants.” PLoS One 8(7): e68098.

Song, L., V. Mishra, M. Ouyang, Q. Peng, M. Slinger, S. Liu and H. Huang (2017). “Human Fetal Brain Connectome: Structural Network Development from Middle Fetal Stage to Birth.” Front Neurosci 11: 561.

Takahashi, E., R. D. Folkerth, A. M. Galaburda and P. E. Grant (2012). “Emerging cerebral connectivity in the human fetal brain: an MR tractography study.” Cereb Cortex 22(2): 455–464.

Turk, E., M. I. van den Heuvel, M. J. Benders, R. de Heus, A. Franx, J. H. Manning, J. L. Hect, E. Hernandez-Andrade, S. S. Hassan, R. Romero, R. S. Kahn, M. E. Thomason and M. P. van den Heuvel (2019). “Functional Connectome of the Fetal Brain.” J Neurosci 39(49): 9716–9724.

Tymofiyeva, O., C. P. Hess, E. Ziv, P. N. Lee, H. C. Glass, D. M. Ferriero, A. J. Barkovich and D. Xu (2013). “A DTI-based template-free cortical connectome study of brain maturation.” PLoS One 8(5): e63310.

Tymofiyeva, O., C. P. Hess, E. Ziv, N. Tian, S. L. Bonifacio, P. S. McQuillen, D. M. Ferriero, A. J. Barkovich and D. Xu (2012). “Towards the “baby connectome”: mapping the structural connectivity of the newborn brain.” PLoS One 7(2): e31029.

Tzourio-Mazoyer, N., B. Landeau, D. Papathanassiou, F. Crivello, O. Etard, N. Delcroix, B. Mazoyer and M. Joliot (2002). “Automated anatomical labeling of activations in SPM using a macroscopic anatomical parcellation of the MNI MRI single-subject brain.” Neuroimage 15(1): 273–289.

van den Heuvel, M. P., S. C. de Lange, A. Zalesky, C. Seguin, B. T. T. Yeo and R. Schmidt (2017). “Proportional thresholding in resting-state fMRI functional connectivity networks and consequences for patient-control connectome studies: Issues and recommendations.” Neuroimage 152: 437–449.

van der Linde, D., E. E. Konings, M. A. Slager, M. Witsenburg, W. A. Helbing, J. J. Takkenberg and J. W. Roos-Hesselink (2011). “Birth prevalence of congenital heart disease worldwide: a systematic review and meta-analysis.” J Am Coll Cardiol 58(21): 2241–2247.

von Rhein, M., A. Buchmann, C. Hagmann, H. Dave, V. Bernet, I. Scheer, W. Knirsch, B. Latal, Heart and G. Brain Research (2015). “Severe Congenital Heart Defects Are Associated with Global Reduction of Neonatal Brain Volumes.” J Pediatr 167(6): 1259–1263 e1251.

Watson, C. G. (2019). brainGraph: Graph Theory Analysis of Brain MRI Data, R package version 2.7.3.

Xia, M., J. Wang and Y. He (2013). “BrainNet Viewer: a network visualization tool for human brain connectomics.” PLoS One 8(7): e68910.

Zalesky, A., A. Fornito and E. T. Bullmore (2010). “Network-based statistic: identifying differences in brain networks.” Neuroimage 53(4): 1197–1207.

Ziv, E., O. Tymofiyeva, D. M. Ferriero, A. J. Barkovich, C. P. Hess and D. Xu (2013). “A machine learning approach to automated structural network analysis: application to neonatal encephalopathy.” PLoS One 8(11): e78824.

## References

Barrat, A., M. Barthelemy, R. Pastor-Satorras and A. Vespignani (2004). “The architecture of complex weighted networks.” Proc Natl Acad Sci U S A 101(11): 3747–3752.

Hernandez, M., G. D. Guerrero, J. M. Cecilia, J. M. Garcia, A. Inuggi, S. Jbabdi, T. E. Behrens and S. N. Sotiropoulos (2013). “Accelerating fibre orientation estimation from diffusion weighted magnetic resonance imaging using GPUs.” PLoS One 8(4): e61892.

Latora, V. and M. Marchiori (2001). “Efficient behavior of small-world networks.” Phys Rev Lett 87(19): 198701.

Latora, V. and M. Marchiori (2003). “Economic small-world behavior in weighted networks.” European Physical Journal B 32(2): 249–263.

Watson, C. G., D. DeMaster and L. Ewing-Cobbs (2019). “Graph theory analysis of DTI tractography in children with traumatic injury.” Neuroimage Clin 21: 101673.

